# Within-species differences in vocal production learning in a songbird are associated with differences in flexible rhythm pattern perception

**DOI:** 10.1101/2022.07.13.499954

**Authors:** Andrew A. Rouse, Aniruddh D. Patel, Samantha Wainapel, Mimi H. Kao

## Abstract

Humans readily recognize a familiar rhythmic pattern, such as isochrony (equal timing between events) across a wide range of rates. This ability reflects a facility with perceiving the relative timing of events, not just absolute interval durations. Several lines of evidence suggest that this ability is supported by precise temporal predictions that arise from forebrain auditory-motor interactions. We have shown previously that male zebra finches, which possess specialized auditory-motor networks and communicate with rhythmically patterned sequences, share our ability to recognize isochrony independent of rate. To test the hypothesis that flexible rhythm pattern perception is linked to vocal learning, we ask whether female zebra finches, which do not learn to sing, can also recognize global temporal patterns. We find that non-singing females can flexibly recognize isochrony but perform slightly worse than males on average. These findings are consistent with recent work showing that while females have reduced forebrain song regions, the overall network connectivity of vocal premotor regions is similar to that in males and supports predictions of upcoming events. Comparative studies of male and female songbirds thus offer an opportunity to study how individual differences in auditory-motor connectivity influence perception of relative timing, a hallmark of human music perception.

## Introduction

The ability to recognize auditory rhythms is critical for many species (Clemens et al., 2021; Garcia et al., 2020; Mathevon et al., 2017), but the underlying neural mechanisms are only beginning to be understood. One area of progress in elucidating the neural circuits for rhythm perception is understanding how communication signals are recognized based on tempo. For example, female field crickets are attracted by male calling songs composed of trains of short sound pulses when the tempo is ∼ 30 syllables/sec. This selectivity is hard-wired and mediated by a small network of interneurons that processes instantaneous pulse rate (Schöneich et al., 2015). While this preference is genetically fixed, in other animals, experience can sculpt neural responses to behaviorally salient call rates. For example, in the mouse auditory cortex, excitatory cells are innately sensitive to the most common pup distress call rate (∼5 syllables/s), but their tuning can broaden to a wider range of rates following co-housing with pups producing calls across a range of rates (Schiavo et al., 2020).

Much less is known about how the brain recognizes rhythmic *patterns* independently of rate. While humans can encode and remember the rate of auditory sequences (Levitin & Cook, 1996), we also readily recognize a given rhythmic pattern, such as isochrony, or equal timing between events, across a broad range of rates (Espinoza-Monroy & de Lafuente, 2021). The ability to recognize a specific rhythmic pattern, whether it is played fast or slow, is present in infants (Trehub & Thorpe, 1989), and is based on recognition of the relative timing of intervals more than on their absolute durations. In addition, the ability to detect and predict isochrony across a range of rates is central to music cognition (Merker et al., 2009; Patel, 2006) and to the positive effects of music-based therapies on a variety of neurological disorders, including normalizing gait in Parkinson’s disease (Dalla Bella et al., 2017; Krotinger & Loui, 2021)

In humans, there is growing evidence that the neural mechanisms underlying perception of relative timing are distinct from those involved in encoding absolute timing (Breska & Ivry, 2018; Grube et al., 2010; Teki et al., 2011). In addition, neuroimaging studies have shown that both auditory *and* motor regions are active when people listen to rhythms, even in the absence of overt movement. Responses in several motor regions are greater when the stimulus has a strong, periodic pulse, or “beat” (Chen et al., 2008; Grahn & Brett, 2007), and transient manipulation of auditory-motor connections using transcranial magnetic stimulation can disrupt beat perception without affecting single-interval timing (Ross et al., 2018). Based on such findings, we and others have suggested that perception of temporal regularity (independent of rate) depends on the interaction of motor and auditory regions: the motor planning system uses information from the auditory system to make predictions about the timing of upcoming events and communicates these predictions back to auditory regions via reciprocal connections (Cannon & Patel, 2021; Patel & Iversen, 2014). Such predictions could support detection of rhythmic patterns independent of tempo because the relative duration of adjacent intervals remains the same across different rates (e.g., 1:1 for an isochronous pattern).

Given that vocal learning species often communicate using rhythmically patterned sequences (Norton & Scharff, 2016; Roeske et al., 2020) and have evolved specialized motor planning regions that are reciprocally connected to auditory forebrain regions, we have hypothesized that vocal learners are advantaged in flexible auditory rhythm pattern perception (Rouse et al., 2021). Consistent with this hypothesis, prior work has shown that zebra finches (*Taeniopygia guttata*) and starlings (*Sternus vulgaris*), two species of vocal learning songbirds, can learn to discriminate isochronous from arrhythmic sound sequences and can generalize this ability to stimuli at novel tempi, including rates distant from the training tempi (Hulse et al., 1984; Rouse et al., 2021, but see ten Cate et al., 2016; van der Aa et al., 2015 for evidence of more limited generalization in zebra finches). This ability, also seen in humans (Espinoza-Monroy & de Lafuente, 2021), demonstrates a facility with recognizing a rhythm based on global temporal patterns, since absolute durations of intervals differ markedly at distant tempi. These findings contrast with similar research conducted with vocal non-learning species. For example, pigeons (*Columbia livia*) can learn to discriminate sound sequences based on tempo but cannot learn to discriminate isochronous from arrhythmic sound patterns (Hagmann & Cook, 2010). Rats (*Rattus norvegicus*) can be trained to discriminate isochronous from arrhythmic sound sequences, but when tested at novel tempi, they show limited generalization, suggesting a strong reliance on absolute timing for rhythm perception (Celma-Miralles & Toro, 2020).

Here, we further test the hypothesis that differences in vocal learning abilities correlate with differences in flexible rhythm pattern perception by taking advantage of sex differences in zebra finches (Nottebohm & Arnold, 1976). Although auditory sensitivity is similar across sex in this species (Yeh et al., 2022), only males learn to imitate song, and the neural circuitry subserving vocal learning is greatly reduced in females. Thus we predicted that male zebra finches would exhibit faster learning rates for discriminating isochronous versus arrhythmic stimuli and/or a greater degree of generalization in recognizing these categories at novel tempi. By using the same apparatus, stimuli, and methods, we can meaningfully compare the flexibility of male vs. female rhythmic pattern perception in this species.

Several prior findings, however, support the opposite prediction - either no sex differences in rhythmic pattern perception or better rhythmic pattern perception in female zebra finches. One reason females may not be disadvantaged in our task is that they analyze male song when choosing a mate, and a previous meta-analysis found that female zebra finches are faster than males at learning to discriminate spectro-temporally complex auditory stimuli (Kriengwatana et al., 2016). In addition, while the volume of vocal motor regions that subserve song performance is substantially greater in male zebra finches, a recent anatomical study found male-typical patterns of connectivity in the vocal premotor region in females, including minimal sex differences in afferent auditory and other inputs (Shaughnessy et al., 2019). Moreover, a recent study of antiphonal calling suggests that female zebra finches may perform better than males on rhythmic pattern processing (Benichov et al., 2016). In that study, both male and female zebra finches could predict the timing of calls of a rhythmically calling vocal partner, allowing them to adjust the timing of their own answers to avoid overlap. This ability to predictively adjust call timing was enhanced in females and was disrupted by lesions of vocal motor forebrain regions, suggesting that brain regions associated with singing in males may subserve auditory perception and/or timing abilities in female zebra finches.

## Methods and analysis

### Subjects

Subjects were 22 experimentally naïve female zebra finches from our breeding colony (mean age=72 ± 8 (SD) days post-hatch (dph) at the start of training; range=61-87 dph). Thirteen females were tested on rhythm discrimination; six females were tested on pitch discrimination and tempo discrimination (see below); and three females did not learn to use the apparatus. For rhythm discrimination, data from 14 age-matched male zebra finches were collected by Rouse et al. (2021). Data from 6/13 females were collected during the same period (March-December 2019) as data for the males; data from the other 7 females were collected following a pandemic-related research hiatus (June-December 2020). All procedures were approved by the Institutional Animal Care and Use Committee.

### Auditory Stimuli

All stimuli used in this experiment were the same as those described in Rouse et al. (2021). Briefly, we used sequences of natural sounds from unfamiliar male conspecifics: either an introductory element that is typically repeated at the start of a song, or a short harmonic stack (Zann, 1996) (see Fig. S1; note differences in amplitude across sequences comprised of different sound elements in Fig. S1B). For each sound, an isochronous sequence (∼2.3 s long) with equal time intervals between event onsets and an arrhythmic sequence with a unique temporal pattern (but the same mean inter-onset interval (IOI)) were generated at two base tempi: 120 ms and 180 ms IOI (Fig. 1A). These tempi were chosen based on the average syllable rate in zebra finch song (∼7-9 syllables per second, or 111-142 ms IOI (Zann, 1996)).

**Figure 1.**
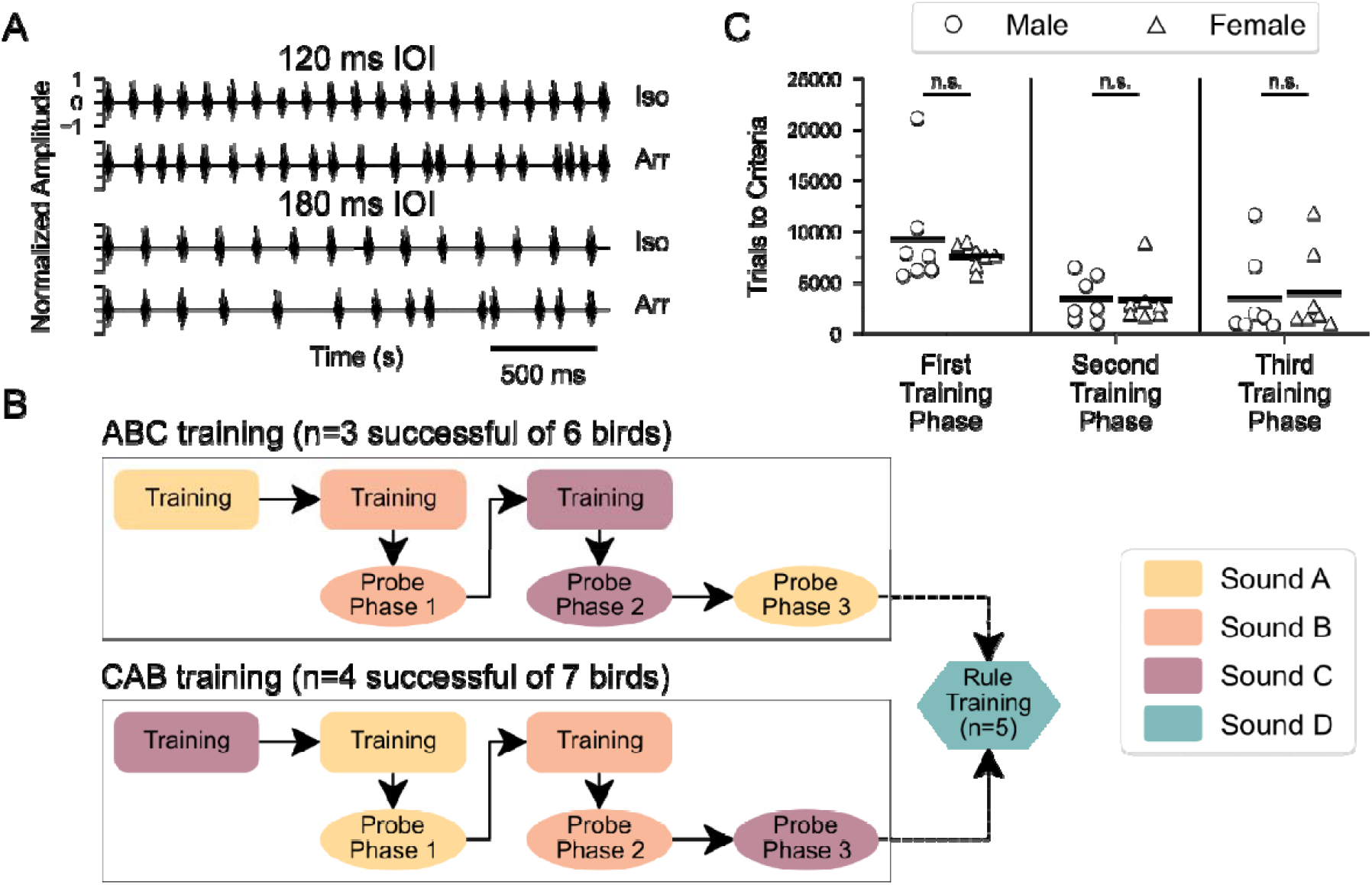
Experiment design for testing the ability to flexibly perceive rhythmic patterns. *(A)* Normalized amplitude waveforms of isochronous (*Iso*) and arrhythmic (*Arr*) sequences of a repeated song element (sound B; see Fig. S1) with 120 ms (*top*) and 180 ms (*bottom*) mean inter-onset interval (IOI). *(B)* Schematic of the protocol. After a pre-training procedure (not shown), birds learned to discriminate between isochronous and arrhythmic sound sequences at 120 and 180 ms IOI, for sounds A, B, and C (‘ABC training’, n=6 females) or starting with sound C followed by A and B (‘CAB training’, n=7 females) (color indicates sound type; see Fig. S1). To test for the ability to generalize the discrimination to new tempi, probe stimuli (144 ms IOI) were introduced after birds had successfully completed two training phases. A subset of birds was then tested with a broader stimulus set using a novel sound element (sound D) and every integer rate between 75 and 275 ms IOI (“Rule Training”; see *Materials and Methods* for details). Color conventions indicating sound type are used in all subsequent figures. *(C)* Comparison of training time, plotted as Trials to Criteria, for male and female birds that completed rhythm discrimination training for all three phases. Symbols denote sex. For this and subsequent figures, data from male birds were collected by Rouse et al. (2021), and are replotted here for comparison. A Mann-Whitney test for each training phase showed no sex-based differences in training time for any phase (*p*>0.05).

Arrhythmic stimuli were generated in an iterative process using MATLAB. Each isochronous stimulus had a known number of intervals, and an equal number of intervals were randomly drawn from a uniform distribution ranging between 0 ms (the minimum possible gap) and 1.5 times the IOI of the corresponding isochronous stimulus (e.g., max IOI of 180 ms and 270 ms for arrhythmic sequences whose mean tempo were 120 and 180 ms IOI, respectively). The process was repeated until it generated a set of intervals that had the same number of intervals, average IOI, and overall duration as the paired isochronous stimulus. To ensure that the arrhythmic pattern was significantly different from isochrony, we also required a minimum SD for interval durations (SD ≥ 10 ms). An additional pair of probe stimuli (one isochronous, one arrhythmic) was generated at a tempo (144 ms IOI) 20% faster/slower than the training stimuli, using the same minimum SD for interval durations and same minimum and maximum IOI constraints.

For “rule training” (see below), an isochronous/arrhythmic pair was generated with a novel harmonic stack (sound D, see Fig. S1A) at tempi ranging from 75 ms to 275 ms IOI in 1 ms steps (i.e., at 201 rates). Each arrhythmic stimulus had a temporally unique pattern. For IOIs < 90ms, the minimum SD of IOI required for arrhythmic sequences was reduced from 10 ms to 1 ms because sound element D was ∼75 ms long (Fig. S1A), making the silent gaps very short and reducing variability in the IOIs. For a subset of these stimulus pairs (n=15/201), the amplitude was inadvertently higher (Fig. S1B).

### Auditory Operant Training Procedure

Training and probe testing used a go/interrupt paradigm described in Rouse et al. (2021). Briefly, female birds were housed singly in a cage that had a trial switch to activate playback of an auditory stimulus and a response switch with a waterspout. Birds were mildly water restricted and worked for water rewards, routinely performing ∼520 trials/day. Pecking the trial switch initiated a trial and triggered playback of a stimulus (∼2.3 s): either rewarded (S+, 50% chance) or unrewarded (S-, 50% chance). For rhythm discrimination experiments, the isochronous patterns were rewarded (S+ stimuli). Trial and response switches were activated 500 ms after stimulus onset, and pecking either switch immediately stopped playback. “Hits” were correct pecks of the response switch during S+ trials. “False alarms” were pecks of the response switch on S-trials and resulted in lights out (up to 25 s) (Gess et al., 2011). If neither switch was pecked within 5 s after stimulus offset, the trial would end (‘no response’). No response to the S-stimulus was counted as a “correct rejection”, and no response to the S+ stimulus was considered a “miss”. During the response window, a bird could also peck the trial switch again to “interrupt” and immediately stop the current trial (and any playback), which was counted as a ‘miss’ or ‘correct rejection’ depending on whether a S+ or S-stimulus had been presented. The use of the trial switch to interrupt trials varied widely among birds and was not analyzed further. Regardless of response, birds had to wait 100 ms after the stimulus stopped playing before a new trial could be initiated.

#### Shaping and performance criteria

To learn the go/interrupt procedure, birds were first trained to distinguish between two unfamiliar conspecific songs (∼2.4 s long, “shaping” phase), one acting as the S+ (rewarded or ‘go’) stimulus, and the other as the S- (unrewarded or ‘no-go’) stimulus. Lights-out punishment was not implemented until a bird initiated ≥ 100 trials. The criterion for advancing to the next phase was ≥ 60% hits, ≥ 60% correct rejections, and ≥ 75% overall correct for two of three consecutive days (Nagel et al., 2010; van der Aa et al., 2015). In each phase, birds were required to reach the performance criterion within 30 days. If a bird failed to reach performance criterion within 30 days in any phase, they were removed from further training/testing. The mean number of trials to criterion for the shaping stimuli was 1734 ± 727 (SD). Three females did not regularly trigger playback of auditory stimuli and were removed from the study without completing shaping.

#### Training: rhythm discrimination

Once a bird reached criterion performance on the shaping stimuli, she was trained to discriminate isochronous versus arrhythmic stimuli (n=13 birds). As described previously, each bird was trained using multiple sound types and multiple stimulus rates. In the first training phase, each bird learned to discriminate two isochronous stimuli (120 ms and 180 ms IOI) from two arrhythmic stimuli (matched for each mean IOI). Once performance criterion was reached, the bird was presented with a new set of stimuli at the same tempi but with a novel sound element (and a novel irregular temporal pattern at each tempo; see Fig. 1B). One group (‘ABC’; n=6) was trained to discriminate sound A stimuli, followed by sound B and then sound C; a second group (‘CAB’; n=7) was trained with sound C, followed by sounds A and B. One female ‘ABC’ bird was presented with stimuli at 3 additional tempi, but those trials were not reinforced and were excluded from analysis.

#### Probe testing/generalization

To test whether birds could generalize the isochronous vs. arrhythmic classification at a novel tempo, females were tested with probe stimuli at 144 ms IOI, 20% slower/faster than the training stimuli (120 ms and 180 ms IOI). Male zebra finches have been previously shown to discriminate between two stimuli that differ in tempo by 20% (Rouse et al., 2021), and we confirmed this in female zebra finches (see below and Fig. S2B). Probe sounds were introduced after a bird had successfully completed two phases of training (see Fig. 1B). Prior to probe testing, the reinforcement rate for training stimuli was reduced to 80% for at least two days. During probe testing, training stimuli (90% of trials) and probe stimuli (10% of trials) were interleaved randomly. Probe trials and 10% of the interleaved training stimuli were not reinforced or punished (Rouse et al., 2021; van der Aa et al., 2015).

#### Rule training

Following probe testing with all three sound types, subjects (n=5 birds) were presented with a new set of isochronous and arrhythmic stimuli using a novel sound (sound D). In this phase, stimulus tempi included every integer rate from 75 ms to 275 ms IOI (201 total rates). Arrhythmic stimuli were again generated independently so that each rate had a novel, random irregular pattern, with the same mean IOI as its corresponding isochronous pattern. Trials were randomly drawn with replacement from this set of 402 stimuli, and all responses were rewarded or punished as during training. This large stimulus set made it unlikely that subjects could memorize individual temporal patterns.

#### Discrimination of other acoustic features

To determine whether the 20% difference between the training and probe stimuli could be detected by female zebra finches, we tested a separate cohort of females on tempo discrimination (n=6; 71 ± 9.1 (SD) dph at start of shaping). Following shaping, these birds were trained to discriminate isochronous sequences of sound A based on rate: 120 ms vs. 144 ms IOI. The rewarded (S+) stimulus was 120 ms IOI for four females and 144 ms IOI for two females. To assess the perceptual abilities of female zebra finches more generally, these females were also tested on their ability to discriminate spectral features using frequency-shifted isochronous sequences of sound B: one shifted 3 semitones up, and the other shifted 6 semitones up (“Change pitch” in Audacity v 2.1.2). Four females were tested with the 6-semitone shift as the S+ stimulus, and two females were tested with the 3-semitone shift as the S+ stimulus.

### Data Analysis

All statistical tests, except for the binomial tests for training and generalization and non-parametric tests for comparisons across perceptual discrimination tasks, were performed in R (v. 3.6.2) within RStudio (v. 1.2.5033). Binomial logistic regressions were performed with lme4 (glmer) statistical package for R, and binomial and non-parametric tests were performed using Python (v3.7.6).

#### Training and generalization testing

To quantify performance, the proportion of correct responses ((Hits + Correct Rejections) / Total number of trials) was computed for each stimulus pair (isochronous and arrhythmic patterns of a given sound at a particular tempo). For training phases, proportion correct was always computed based on the last 500 trials. For probe testing, performance was computed for 80 probe trials. The proportion of correct responses was compared to chance performance (*p*=0.5) with a binomial test using α=0.05/2 in the training conditions (2 tempi), α=0.05/3 for probe trials (3 tempi). A linear least-squares regression was used to examine the correlation between each subject’s average performance on interleaved training stimuli and probe stimuli.

To take into account incorrect responses to the S^−^ stimuli, we also calculated the d-prime measure for both training stimuli and probe stimuli:

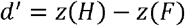

where *H* is the proportion of correct responses to the S+ stimulus (Hits) and *F* is the proportion of incorrect responses to the S-stimulus (false alarm rate). A d’ value of 0 is equivalent to chance performance, and d’ values greater than 1 are considered ‘good’ discrimination.

Performance of successful females (n=7/13) during probe trials was compared to that of successful male zebra finches (n=7/10) collected in a prior study (Rouse et al., 2021). Performance across all probe trials (n=240 probe trials, 80 probe trials per song element x 3 song elements) was analyzed with a binomial logistic regression using a generalized linear mixed model with sex, probe phase number, and sex-by-probe phase number interaction as fixed effects and subject as a random effect.

#### Rule training

As described previously in Rouse et al. (2021), the first 1000 trials were analyzed for each bird (n=5) to minimize any potential effect of memorization. Trials were binned by IOI in 10 ms increments, and the number of correct responses was analyzed with a binomial logistic regression using a generalized linear mixed model (tempo bin as a fixed effect and subject as a random effect). Performance in each bin was compared to performance in the 75-85 ms IOI bin, where performance fell to chance since the degree of temporal variation in inter-element intervals was severely limited by the duration of the sound element. To identify possible sex differences, we performed an additional analysis using data from a range of tempi (95-215 ms) in which both males and females performed significantly above chance (p<0.005; binomial test). The number of correct and incorrect trials were calculated per sex and analyzed for significant group differences with a χ2 test.

#### Discrimination of other acoustic features

Performance on tempo or frequency discrimination was analyzed in the same manner as for rhythm discrimination: the proportion of correct responses in the last 500 trials was compared to chance performance using a binomial test with α=0.05.

#### Reaction Time

Given that our task involves perception of temporal patterns, we quantified how long birds listened to the stimuli before responding. To estimate the duration of the stimulus that a bird heard prior to responding, the median time between trial initiation and response selection (when the bird pecked a switch) was computed for the last 500 trials in a training phase. For the rhythm discrimination birds, only the first training phase (prior to probe testing) was used. Median reaction time values were compared between 3 groups: 14 birds that completed all rhythmic training phases, 9 birds that failed to complete rhythm discrimination training, and 10 birds that learned the frequency discrimination (data from 6 females in this study and from 4 males collected as part of Rouse et al. (2021)). No sex differences were observed when pooling reaction time data across rhythm discrimination and frequency discrimination tasks (Mann-Whitney U test: U=0.69; *p* = 0.49). A Kruskal-Wallis test was used to compare differences across groups, followed by a Dunn’s post-hoc test with Bonferroni correction for multiple comparisons.

## Results

### Rhythmic pattern training and generalization testing in female zebra finches

To test the ability of female songbirds to recognize a rhythmic pattern based on the relative timing of events, 13 female zebra finches were first trained to discriminate isochronous from arrhythmic sequences using a go/no-go paradigm with three training phases (Figs. 1A–B & S1). Seven out of 13 females learned to discriminate these sequences in all three phases, each of which used a different zebra finch song element. The number of trials for these females to reach criteria for rhythmic discrimination (see Materials and Methods) was comparable to that of male zebra finches tested previously (Fig. 1C; data from male birds in Rouse et al. (2021)). Fig. 2A shows the time course of learning for a representative female zebra finch (*y7o97*), which gradually learned to withhold her response to the arrhythmic stimulus. Across the seven successful females, rhythm discrimination performance was ∼81% accurate (mean d’ = 1.9) at the end of the first training phase (Fig. 2B, left column), and a comparable accuracy level was attained in each training phase (median = 80% proportion correct; mean d’ = 1.7 see Fig. S2A for learning curves for each bird on the rhythm discrimination task).

**Figure 2.**
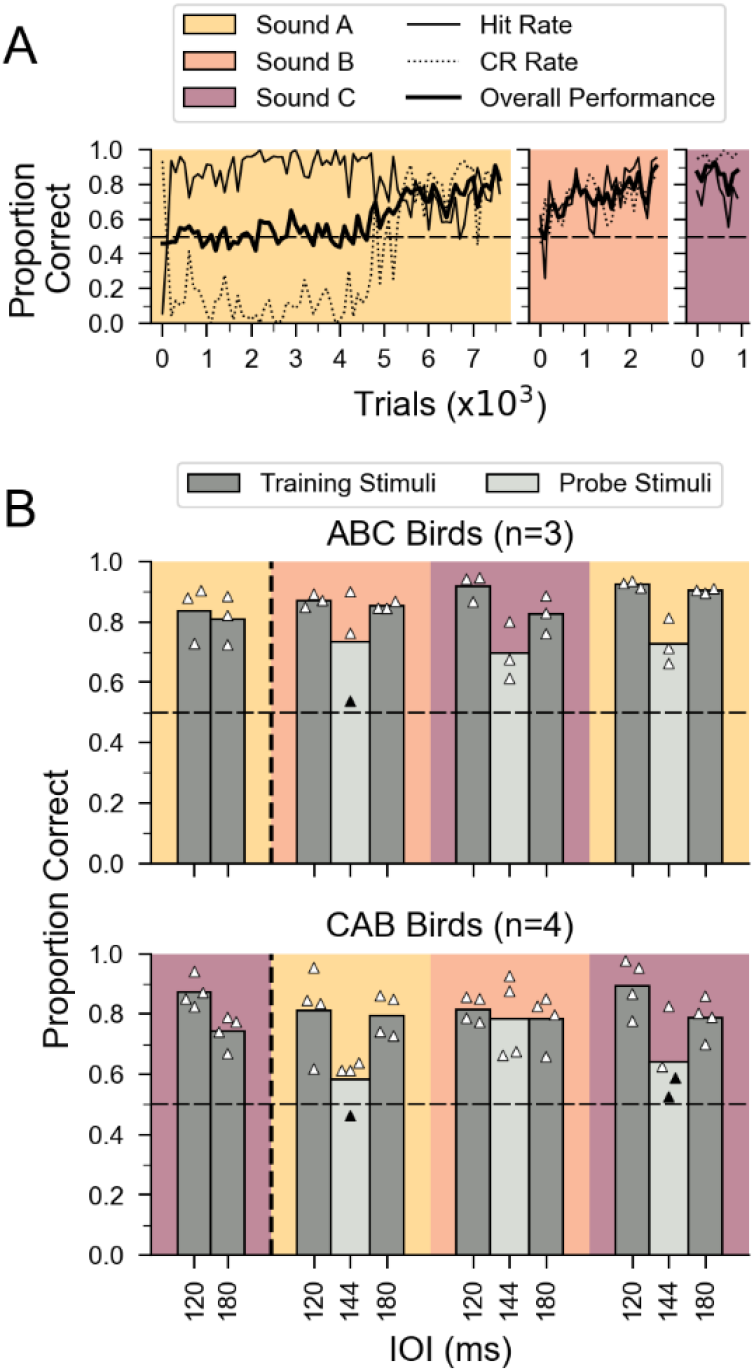
Learning of rhythmic pattern discrimination and generalization to new tempi for female birds. (*A)* Learning curves for rhythm discrimination for a representative female bird (*y7o97*) across three training phases (ABC training) in 100-trial bins. Overall proportion of correct responses is shown by *thick black* line. Proportion correct for isochronous stimuli (S+, “hit rate*”*) is indicated by *thin black* line; proportion correct for arrhythmic stimuli (S-, “correct rejection*”*) is indicated by *dotted line*. Chance performance is indicated by dashed horizontal line. Data are plotted until criterion performance was reached. (*B*) Results for rhythm discrimination training and probe testing for successful female birds (n=7). Data left of the vertical dashed line show performance in the final 500 trials of the first rhythm discrimination training phase (no probe testing, see Fig. 1B). Data right of the vertical dashed line show performance during probe testing with stimuli at an untrained tempo of 144 ms IOI (*light gray*) and for interleaved training stimuli (*dark gray*). Triangles denote performance for each female (n = 3 probe tests/bird x 7 birds). All seven females were able to generalize the isochronous versus arrhythmic discrimination to new tempi (20% different from the training tempi) for at least two sounds. Performance on probe stimuli was significantly above chance for 17 out of 21 probe tests (*white* triangles; *p<*.0.0167, binomial test with Bonferroni correction; black triangles denote performance not significantly different from chance). Average performance across birds in each group is indicated by bars.

After successful completion of two phases of rhythm discrimination, females were then asked whether they could generalize the discrimination of isochronous versus arrhythmic stimuli at a novel tempo (Fig. 1B, ‘probe testing’). Fig. 2B shows the performance on randomly interleaved training stimuli (*dark gray bars*; 90% of the trials) and 80 probe stimuli at a novel tempo (*light gray bars;* 10% of the trials) for the seven females that successfully completed all three phases of rhythm discrimination training. Performance on probe stimuli was significantly above chance for 17 of 21 probe tests (n=3 probe tests/bird x 7 birds, *p*<0.0167, binomial test with Bonferroni correction), indicating that female zebra finches robustly generalized the discrimination of isochronous vs. arrhythmic stimuli to a novel tempo distant from the training tempi (Fig. 2B).

### Sex differences in rhythm pattern perception

To test the hypothesis that the ability to flexibly perceive rhythmic patterns is linked to vocal learning, we directly compared performance of female zebra finches, which do not learn to imitate song, with that of vocal learning males (Fig. 3 & Fig. S3). Across all generalization tests, discrimination of isochronous versus arrhythmic stimuli at a novel tempo (probe stimuli) was significantly lower for female zebra finches compared to males (n=7 males and 7 females; 240 probe trials per bird; mixed-effects logistic regression, *p* = 0.033). The observed sex difference in the ability to generalize rhythm discrimination did not reflect differences in motivation to perform the task. On average, female zebra finches performed ∼520 trials/day, compared to ∼530 trials/day for males in a prior study. Across all subjects (n=14), performance on interleaved training stimuli (learned discrimination) was positively correlated with performance on probe stimuli (measure of generalization; Fig. S4; *p* = .0375).

**Figure 3.**
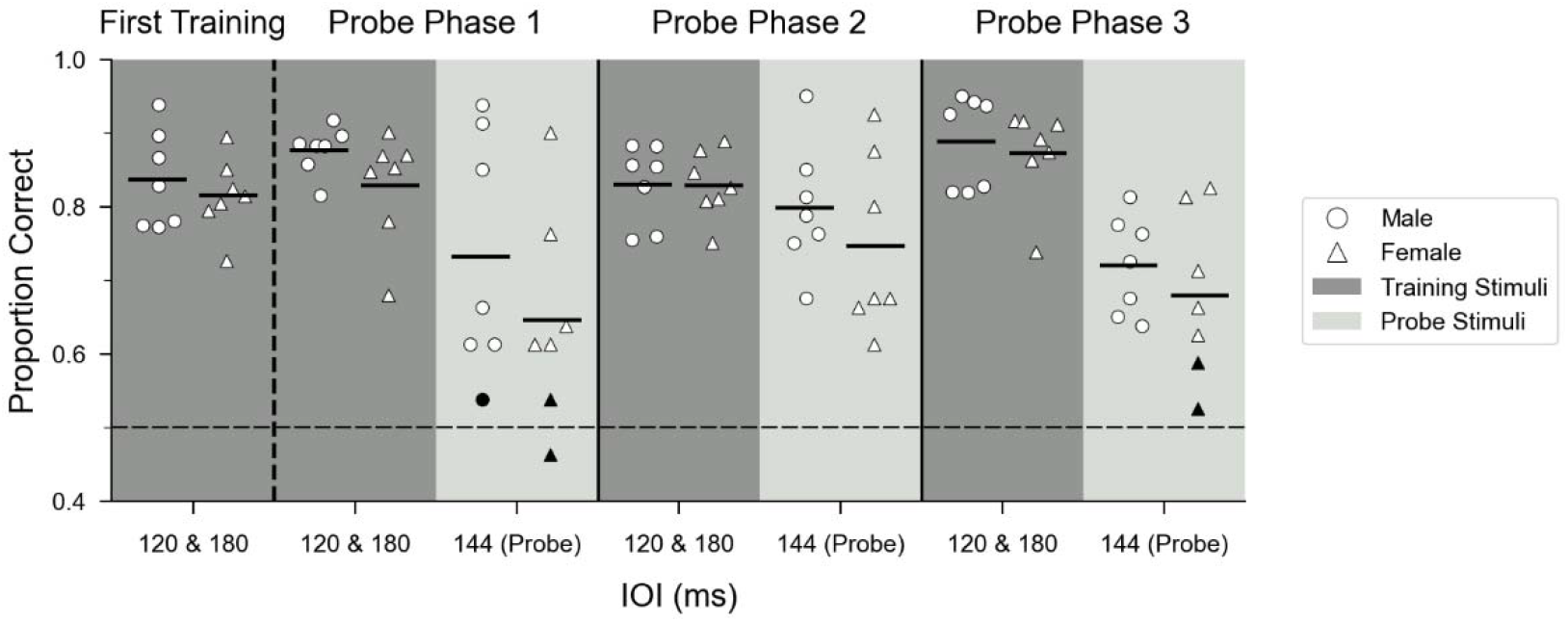
Comparison of generalization of rhythm discrimination by sex. Mean performance of all successful males (*circles;* n=7) and females (*triangles*, n=7) during the last 500 trials of the first training phase (first column) and for probe (*light gray*) and interleaved training stimuli (*dark gray)* in subsequent phases. Data are collapsed across training order, and interleaved training trials are combined across tempo. Dashed horizontal line indicates chance performance. Filled black symbols are not significantly different from chance. Across probe tests, average performance was significantly lower for females than for males by ∼9% (*p<*0.05, mixed-effects logistic regression).

As a second test of the ability to recognize temporal regularity independent of tempo, five females that completed rhythm discrimination training were tested with sequences of a new sound element (sound D, see Figs. S1-S2) across a wide range of tempi (75-275 ms IOI; “rule training”). For each trial, one of 402 stimuli was played (with equal odds of an isochronous or arrhythmic sequence), reducing the likelihood that correct discrimination was based on memorization. Fig. 4 shows the average performance of all birds over the first 1000 trials, broken into 10 ms IOI bins (∼50 trials per bin). For female birds, performance was best between 95 and 215 ms IOI (tempi ∼20% slower to ∼25% faster than the original training range: darker bars with >=2 black asterisks, *p*<0.005; mixed effect logistic regression). Performance fell to chance at faster tempi (75-85 ms IOI), when temporal variability in IOIs was limited by the length of the sound element, and at slower tempi (225-275 ms IOI). To directly compare performance to that of male zebra finches, the number of correct and incorrect trials within the 95-215 ms IOI tempo range (where both sexes performed well above chance) were grouped by sex. In this range, female performance was significantly worse than male performance (χ_2_(1) = 13.919, *p*<0.001).

**Figure 4.**
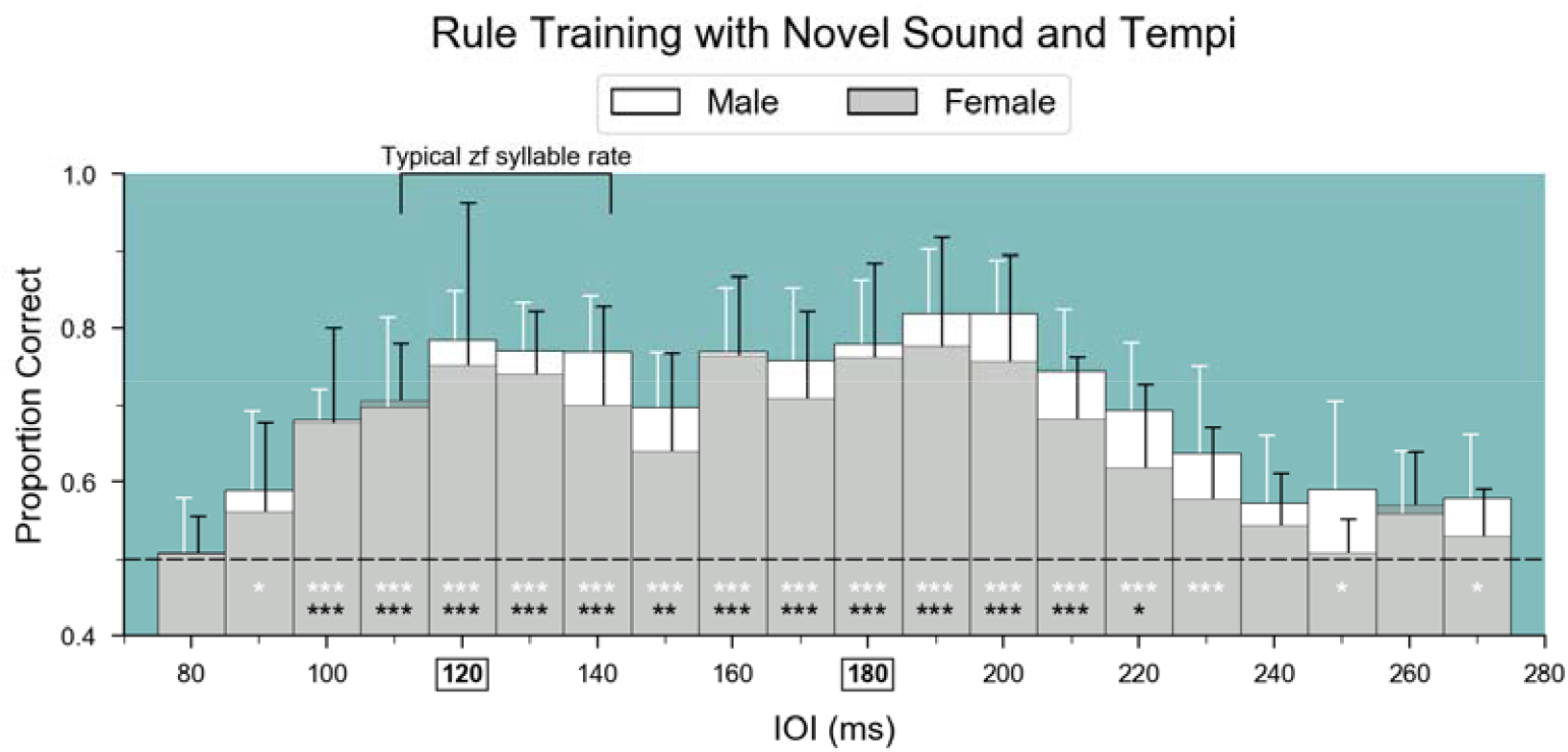
Sex differences in generalization of the learned discrimination across a wide range of tempi. Average performance of 5 female birds (dark bars) during the initial 1000 trials of “Rule training” with a novel sound element (‘D’; color conventions as in Figure 1). Mean performance ± SD is plotted for 10 ms IOI bins. Chance performance is indicated by the horizontal dashed line. Female performance for binned IOIs from 95-225 ms is significantly above performance in the 75-85 ms IOI bin, where performance was at chance (\*\*\**p<*0.001; \*\**p<*0.005; \**p<*0.05, mixed-effect logistic regression, black asterisks). Data from 7 males from Rouse et al. (2021) are plotted for comparison (white bars and asterisks). IOIs used in rhythm discrimination training phases before rule training are shown in bold and boxes on the x-axis.

### Reaction times for discrimination of acoustic features

Across female birds that succeeded in rhythm discrimination training (n=7), the average reaction time at the end of the first training phase was ∼1.20 s after stimulus onset, or ∼50% of the duration of the stimulus train (Fig. 5 & Table S1). Data from previously tested males that succeeded in rhythm discrimination training (n=7) shows a similar average reaction time (1.30 s), indicating that both males and females heard ∼7–11 intervals on average before responding. In contrast, the average reaction time for the birds that did not reach the criterion for rhythm discrimination during training was ∼1.0 s for females (n=6) and 0.9 s for males (n=3).

**Figure 5.**
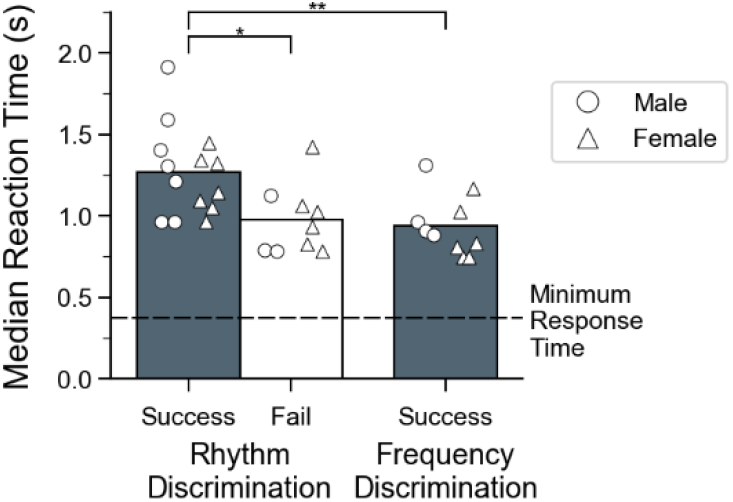
Reaction times during rhythmic pattern discrimination and frequency discrimination tasks. Median reaction times for the final 500 responses in the first training phase are shown for three groups of birds. For rhythm discrimination, data are shown for birds which completed all three training phases (first bar, “success”, n=7 males and 7 females) and birds that did not complete rhythm discrimination training (middle bar, “fail”, n=3 males and 6 females). A separate cohort of birds all succeeded at discriminating isochronous sequences that differed in frequency (third bar, n=4 males and 6 females). The 500ms period between trial start and activation of the response switch is indicated by the horizontal dashed line. Symbols indicate the median reaction time for each bird; bars indicate means of these values collapsed across sex. A Kruskal-Wallis test showed a significant effect of group (H(2) =11.33; p = 0.003). Successful rhythm discrimination birds responded significantly slower than the other two groups (*p*=0.03 compared to the failed birds, *p=*0.008 compared to the frequency discrimination birds; post-hoc Dunn’s test with Bonferroni correction).

Shorter reaction times, however, do not necessarily indicate poor perceptual discrimination. In a separate cohort of birds that learned to discriminate isochronous sequences that differed in frequency by 3 semitones (i.e., a quarter of an octave; n = 6 females; average proportion correct responses in last 500 trials = 91%; mean d’ = 2.8; see Fig. S2B), the average median reaction times for successful discrimination was ∼0.9 s. This suggests that unlike rhythm discrimination, discrimination based on spectral features does not require hearing long stretches of the sequence.

Comparison of reaction times between successful rhythm discrimination birds, unsuccessful rhythm discrimination birds, and successful frequency discrimination birds showed a significant effect of group (Fig. 5 & Table S1; Kruskal-Wallis, H(2)=11.33; *p*=0.003). Dunn’s post-hoc tests showed that the median reaction times of the successful rhythm birds were significantly higher (slower) than both the unsuccessful rhythm birds and the successful frequency birds. Taken together, these results suggest that success at rhythm discrimination may be related to how long birds listened to the stimuli before responding. Indeed, even among the birds that successfully generalized the discrimination of temporal regularity across a wide range of tempi, median reaction time was longer for the tempi where the discrimination was more difficult (i.e., when the proportion of correct responses was closer to chance performance: 75-85 ms IOI and >235 ms IOI; Fig. S5).

## Discussion

To test the hypothesis that differences in the capacity for flexible rhythm pattern perception correlate with differences in vocal learning, we investigated the ability of a sexually dimorphic songbird to recognize a fundamental rhythmic pattern common in music and vocal communication – isochrony (Norton & Scharff, 2016; Ravignani & Madison, 2017; Roeske et al., 2020; Savage et al., 2015). In zebra finches and many other songbirds, only the males learn to sing and possess pronounced forebrain motor regions for vocal production learning (Nottebohm & Arnold, 1976). Male zebra finches have been shown to discriminate isochronous versus arrhythmic patterns in a tempo-flexible manner (Rouse et al., 2021), and we predicted that female zebra finches would perform less well when tested with the same rhythm discrimination and generalization tasks. Using a sequential training paradigm with multiple sound types and tempi, we found that about half of females tested (n=7/13) could learn to discriminate isochronous from arrhythmic patterns in all three phases of our training paradigm (Fig. 1B), and the learning rates for these females were similar to those of males (Fig. 1C). In each training phase, birds were required to reach a performance criterion of >75% correct responses within 30 days, so it is possible that more females may have learned the discrimination with additional training. Females that completed all three phases of training showed generalization of the learned discrimination for a novel tempo 20% different from either training tempo (Fig. 2B). Importantly, zebra finches can detect a 20% tempo difference (Fig. S2B, “tempo discrimination”), suggesting that their ability to recognize isochrony in the probe stimuli did not derive simply from the physical similarity between the training stimuli and the probe stimuli (i.e., generalization around a familiar training stimulus). Notably, birds that succeeded in our rhythm training paradigm listened for longer to stimulus sequences than birds who failed (Fig. 5, left two bars).

We also found that zebra finches that completed training were able to discriminate isochronous from arrhythmic stimuli across a range of tempi far beyond the training tempi and beyond the natural tempo range of zebra finch vocalizations (Fig. 4). In this “rule training” experiment, birds heard an isochronous or arrhythmic pattern randomly drawn from 201 different tempi, with each tempo having a unique arrhythmic pattern. Responses to all stimuli were reinforced but only the first 1000 trials were analyzed (i.e., ∼5 trials/tempo), allowing little time to learn this discrimination based on reinforcement. At very fast tempi, failure to discriminate isochronous from arrhythmic patterns was expected, since the short inter-onset intervals allowed for minimal temporal variation (Rouse et al., 2021, Fig. S3A). The failure to discriminate at slower tempi suggests an inability to accurately predict the timing of the next event in isochronous sequences, as happens with humans listening to very slow metronomes (Bååth et al., 2016). Our finding of flexible rhythm perception contrasts with previous work that reported limited ability of zebra finches to recognize isochrony independent of tempo (ten Cate et al., 2016; van der Aa et al., 2015).

Based on those studies, the authors proposed that zebra finches tend to recognize rhythms based on absolute durations rather than relative timing patterns. While we cannot rule out the possibility that successful discrimination in our rule training experiment was due, in part, to detection of absolute durations and learning to respond to stimuli close to the training stimuli, we believe that several factors in our study encouraged birds to recognize isochrony independent of tempo. First, unlike prior studies, we used multiple training phases, each with long sequences of a different conspecific vocalization (Fig.1A & Fig. S1) and novel arrhythmic temporal patterns. The use of stimuli with task-irrelevant spectral and amplitude differences across training phases as well as multiple training tempi may have promoted attention to the global temporal pattern, which was the only relevant dimension on which rewards were based. Second, while prior studies of rhythm discrimination tested adult zebra finches that had completed sensorimotor learning, we trained and tested younger birds still in the sensorimotor learning phase (mean age = 72 ± 8 (SD) dph at the start of training).

Differences between our findings and earlier work suggest that zebra finches may rely on absolute timing for recognizing rhythmic patterns under some circumstances and on relative timing under different circumstances, depending on variables such as the training method, the stimuli (e.g., conspecific vocalizations vs. human musical sounds), and age. In future work, it would be interesting to explore this further, e.g., by repeating our experiment with adult zebra finches to see if age is an important factor. While female zebra finches can learn to recognize isochrony across a broad range of tempi, we found small but significant differences in performance between males and females, consistent with our hypothesis that differences in vocal learning abilities correlate with differences in flexible rhythm pattern perception. First, across all three tests of generalization at a novel tempo, males outperformed females (Fig. 3; average proportion of correct responses during probe trials: 75% vs. 69%; average d’=1.7 vs. 1.2). Second, males consistently slightly outperformed females when tested with stimuli across a broad range of tempi at which both sexes performed well above chance (Fig. 4; 95-215 ms IOI; average proportion correct: 76% vs 72%; average d’=1.5 vs. 1.2). In addition, males were able to recognize isochrony across a broader range than females (30% faster to 25% slower than the original training stimuli for males; ∼25% faster to 20% for females, Fig. 4). These differences did not reflect differences in motivation to perform the task; on average, males and females performed comparable numbers of trials/day. Finally, the proportion of females that successfully recognized isochrony based on global temporal patterns (n=7/13) was lower than that of males tested previously (n=7/10), although more data would be needed to determine if this difference is reliable.

It is important to note that the sex difference we observed, while consistent with our hypothesis, is an *average* difference. Modest, but consistent, sex differences are well known in biology (e.g., on average, men are taller than women, due, in part, to sex-biased gene expression (Naqvi et al., 2019)), but these differences typically pertain to anatomy, not cognition (Zentner, 2021). Prior work on sex differences in the auditory domain has focused largely on neural mechanisms in the periphery (Berninger, 2007; Gall et al., 2013; Krizman et al., 2020, 2021), although more recent work has demonstrated hormone-mediated differences in forebrain auditory responses to conspecific songs in birds (Brenowitz & Remage-Healey, 2016; Krentzel & Remage-Healey, 2015). Here, we show a small, but consistent sex difference in an auditory cognitive task – flexible rhythm pattern recognition – in a sexually dimorphic bird. While our sample sizes are in line with other studies of sex differences in auditory processing in birds (Benichov et al., 2016; Kriengwatana et al., 2016), additional studies are needed to confirm this difference.

Just as some women are taller than some men, we find that some individual female zebra finches can outperform individual males in our tasks (e.g., probe tests in Fig. 3). How can this be reconciled with our hypothesis of a link between the neural circuitry for vocal learning and flexible rhythm pattern perception? Recent neuroanatomical work found that although female zebra finches possess smaller vocal motor regions, the overall network connectivity of vocal premotor regions is similar in male and female zebra finches (Shaughnessy et al., 2019). Similarities in ascending auditory inputs to premotor regions, in particular, suggest that auditory processing mechanisms may be conserved across sex. This raises the question of whether individual variation in auditory-motor circuitry correlates with individual differences in flexibility of rhythmic pattern perception, irrespective of sex. Indeed, in humans, the strength of cortical auditory-motor connections predicts individual differences in rhythmic abilities (Blecher et al., 2016; Vaquero et al., 2018).

Our finding that female zebra finches perform slightly worse than males in recognizing isochrony contrasts with a prior finding that female zebra finches outperform males in a task involving temporal processing of rhythmic signals (Benichov et al., 2016). In that study, birds (6 males and 6 females) called antiphonally with a robotic partner that emitted calls at a rate of 1 call/second. Birds quickly learned to adjust the timing of their calls in order to avoid a jamming signal introduced at a fixed latency after the robot call, with females showing more pronounced adjustments of call timing than males. While both our study and that study highlight the importance of prediction for rhythm perception, the underlying mechanisms for predicting upcoming events may differ between the two studies. In principle, avoidance of the jamming signal could have resulted from learning a single temporal interval – the time between the robot call and the jamming signal. In contrast, in our task birds had to learn to recognize the relative timing of successive events and to respond to the same pattern, even when absolute interval durations changed. Prior work suggests distinct mechanisms for single-interval versus relative timing (Breska & Ivry, 2018; Grube et al., 2010; Teki et al., 2011), so a male advantage on a task that depends on relative timing is not necessarily inconsistent with a female advantage in single interval timing.

Our work adds to a growing body of research suggesting that sensitivity to rhythmic patterns is widespread across animals. For example, Asokan et al. (2021) found that the responses of neurons in the primary auditory cortex of mice are modulated by the rhythmic structure of a sound sequence. While neurons in the midbrain and thalamus encode local temporal intervals with a short latency, cortical neurons integrate inputs over longer a timescale (∼1 s), and the timing of cortical spikes differs depending on whether consecutive sounds are arranged in a repeating rhythmic pattern or are randomly timed. In Mongolian gerbils (*Meriones unguiculatus*), midbrain neurons have also been shown to exhibit context-dependent responses: on average, neural responses were greater for noise bursts that occurred on the beat of a complex rhythm compared to when the same noise bursts occurred off-beat (Rajendran et al., 2017). Similarly, in rhesus monkeys (*Macaca mulatta*), occasional deviant sounds in auditory sequences elicit a larger auditory mismatch negativity in electroencephalogram recordings when those sequences had isochronous vs. arrhythmic event timing (Honing et al., 2018). While these studies found context-dependent modulation of auditory responses, they did not test the ability of the animals to recognize a learned rhythm independently of tempo. Demonstrating this ability requires behavioral methods, and an important lesson from prior research is that training methods can strongly influence to what extent such abilities are revealed (e.g., compare van der Aa et al. (2015) with Rouse et al. (2021) and Hulse et al. (1984) with Samuels et al. (2021); see also Bouwer et al. (2021)).

Our study investigated rhythm perception in a sexually dimorphic songbird, and a natural question is to what extent similar findings would be obtained in mammals with differing vocal learning capacities, ranging from animals that can modify innate vocalizations in response to sensory cues, such as rodents, bats, and nonhuman primates, to animals that learn to imitate vocalizations, such as seals and cetaceans (Hage et al., 2016; Janik, 2014; Stansbury & Janik, 2019; Takahashi et al., 2017; Vernes & Wilkinson, 2019; Wirthlin et al., 2019). Rats can be trained to discriminate isochronous from arrhythmic sequences, but show limited generalization when tested at novel tempi (Celma-Miralles & Toro, 2020). Macaques can also be trained to discriminate isochronous from arrhythmic sequences, but their ability to do this flexibly at novel tempi distant from training tempi has not been tested (Espinoza-Monroy & de Lafuente, 2021). Interestingly, macaques can be trained flexibly entrain their saccades or tapping movements to isochronous sequences at different tempi (Gámez et al., 2018; Takeya et al., 2017), and population-level neural activity consistent with a relative timing clock has been reported in premotor regions (pre-supplementary motor area and supplementary motor area) during the tapping task (Gámez et al., 2019; Merchant et al., 2015). Thus, they would be an interesting animal model for investigating flexible rhythm pattern perception in the absence of movement. More recently, vocal learning harbor seals (*Phoca vitulina*), have been shown to discriminate isochronous from arrhythmic sequences (Verga et al., 2022), though their ability to do this flexibly across a wide range of tempi remains unknown. Thus, comparative study of flexibility of rhythmic pattern perception is an area ripe for further behavioral and neural investigation. More generally, the ability to relate neural activity to perception and behavior is critical for understanding the contributions of motor regions to detecting temporal periodicity and predicting the timing of upcoming events, two hallmarks of human rhythm processing that are central to music’s positive effect on a variety of neurological disorders.

## Acknowledgments

We thank T. Gardner and his laboratory for equipment and technical assistance with the operant chambers and T. Gentner for sharing the Pyoperant code. We thank members of the Kao lab for useful discussions and comments on earlier versions of this manuscript.

## Supplemental Figures

**Figure S1.**
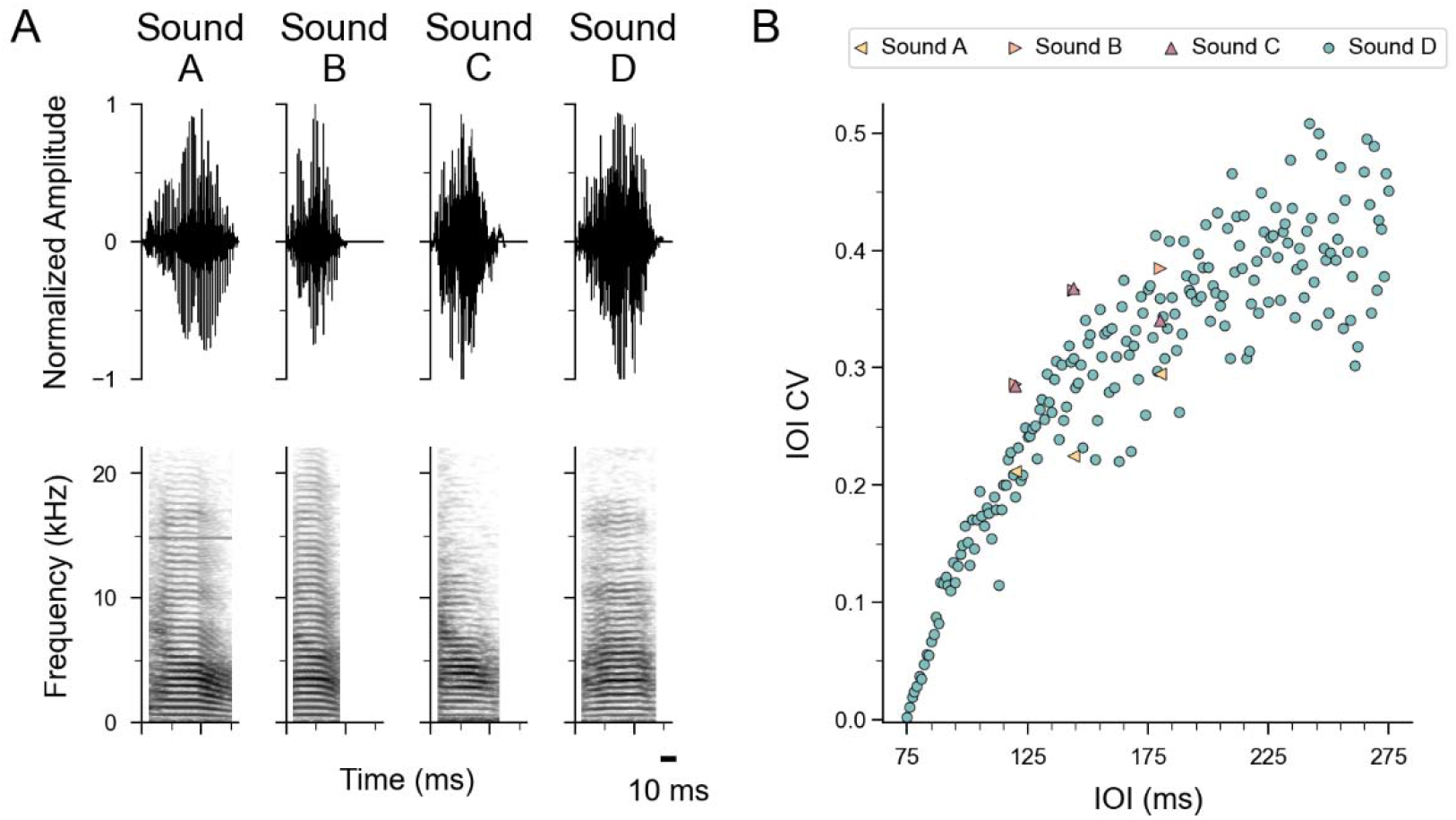
Stimulus waveforms, spectrograms, and CVs. *(A)* Normalized amplitude waveforms (top) and spectrograms (bottom) of the four song elements used in stimulus creation. Durations of the sound elements are as follows: Sound A = 79.9 ms, Sound B = 51.5 ms, Sound C = 63.1 ms, and Sound D = 74.7 ms. *(B)* Coefficient of variation (CV) of inter-onset-intervals for all arrhythmic stimuli against mean stimulus IOI. Note that as the average IOI approaches the length of the source sound element, IOI variability approaches 0. Sound files are available at https://dx.doi.org/10.17632/fw5f2vrf4k.2

**Figure S2.**
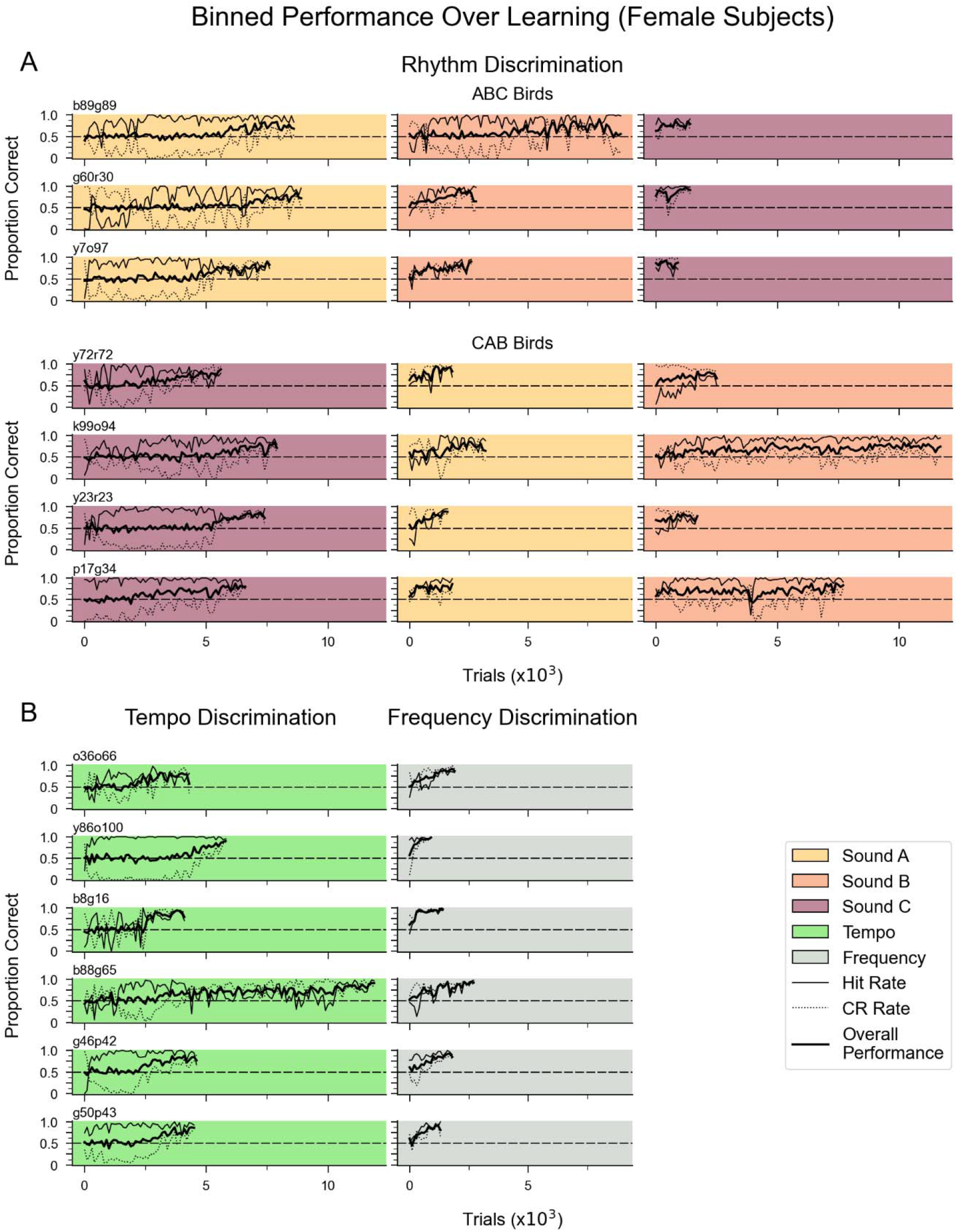
Learning curves for discrimination of stimuli based on rhythmic pattern, tempo, or frequency. (*A*) Learning curves for 7 female birds that successfully discriminated isochronous vs. arrhythmic stimuli within 30 days for each sound type. (*B*) Learning curves for a separate cohort of females (n=6) trained to discriminate isochronous sequences of sound A that differed in tempo (120 ms IOI vs. 144 ms IOI) or frequency (shifted up 3 semitones vs. 6 semitones). All females tested met the criterion for discriminating stimuli based on tempo or frequency. Conventions as in Fig. 2A.

**Figure S3.**
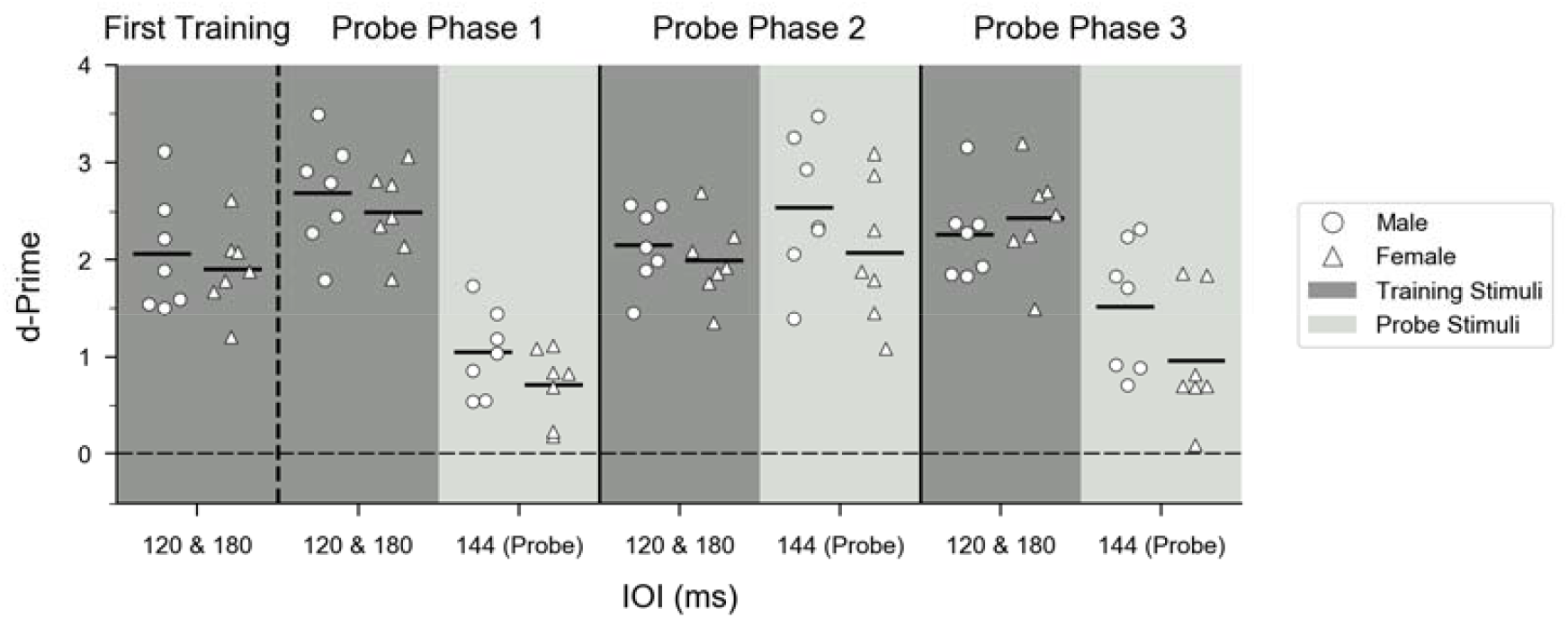
Comparison of generalization of rhythm discrimination by sex. Mean d’ values for all successful males (*circles;* n=7) and females (*triangles*, n=7) during the last 500 trials of the first training phase (first column) and for probe (*light gray*) and interleaved training stimuli (*dark gray)* in subsequent phases. Data are collapsed across training order, and interleaved training trials are combined across tempo. Dashed horizontal line (d’=0) indicates chance performance; d’ =3 is close to perfect performance.

**Figure S4.**
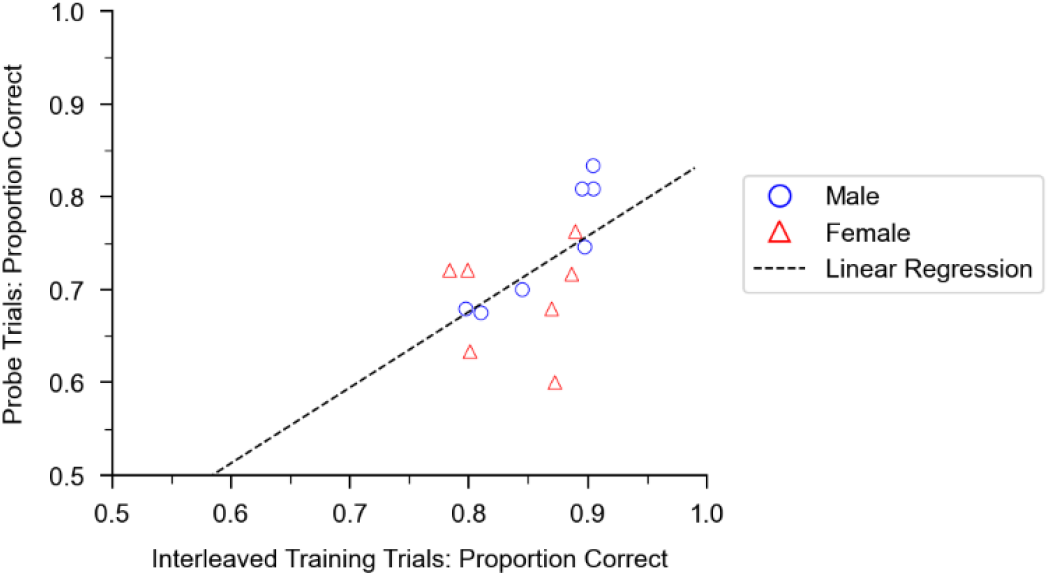
Relationship between performance on probe trials and interleaved training trials. Across birds that successfully completed all three training phases (n=14), there was a significant positive correlation between the accuracy on interleaved training stimuli (learned discrimination) and probe stimuli (generalization); least squares regression fit; R^2^ =0.31; p = 0.0375). For each bird, data were combined across all three sound elements; each data point is the average performance across 240 probe stimuli and ∼2000 interleaved training stimuli.

**Figure S5.**
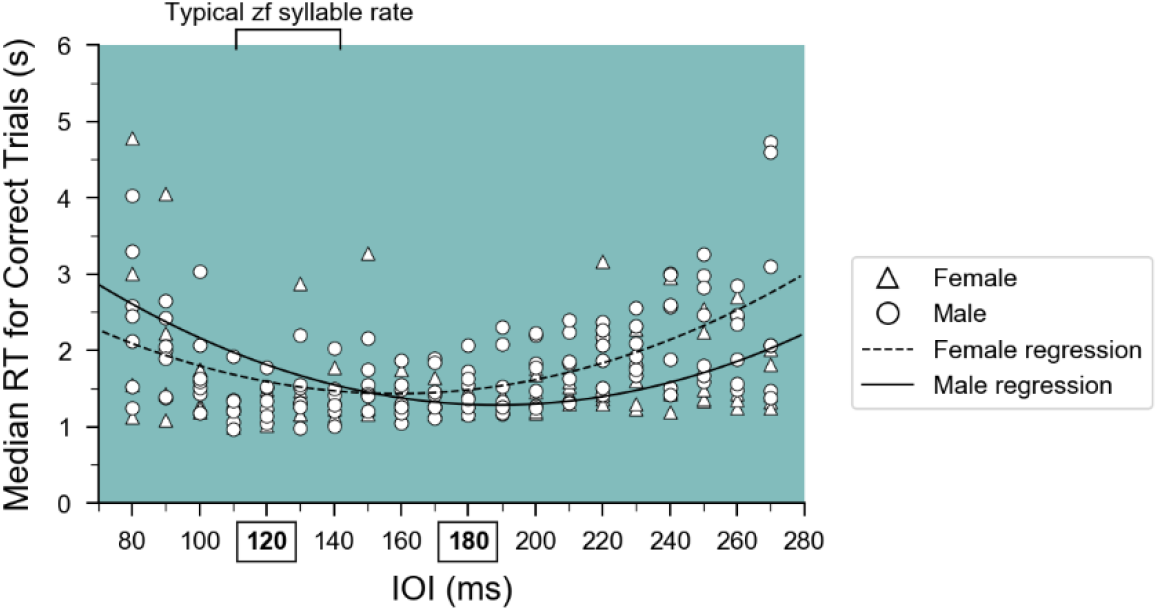
Reaction time (RT) across a broad range of tempi. Median reaction times for correct trials versus tempo during Rule Training for male (*circles*) and female (*triangles*) birds. RT tended to be longer at the end of the tempo range (<100 ms and >220 ms), when discrimination of isochronous vs. arrhythmic stimuli fell to chance. Fitted quadratic curves are plotted to help illustrate the observed relationship.

**Table S1.** Summarized RTs from the last 500 trials of the first training block for each subject. Only trials in which the bird pecked a switch are included.

| Group | Discrimination | Sex | RT median (s) | RT IQR (s) |
| --- | --- | --- | --- | --- |
| Rhythm | Success | Male | 0.965 | 0.440 |
| Rhythm | Success | Male | 1.404 | 1.454 |
| Rhythm | Success | Male | 0.965 | 0.360 |
| Rhythm | Success | Male | 1.915 | 1.288 |
| Rhythm | Success | Male | 1.305 | 0.974 |
| Rhythm | Success | Male | 1.210 | 0.692 |
| Rhythm | Success | Male | 1.590 | 1.473 |
| Rhythm | Success | Female | 1.095 | 0.565 |
| Rhythm | Success | Female | 1.344 | 0.370 |
| Rhythm | Success | Female | 0.965 | 0.660 |
| Rhythm | Success | Female | 1.445 | 0.520 |
| Rhythm | Success | Female | 1.325 | 2.584 |
| Rhythm | Success | Female | 1.050 | 0.376 |
| Rhythm | Success | Female | 1.145 | 0.416 |
| Rhythm | Fail | Male | 0.790 | 0.247 |
| Rhythm | Fail | Male | 0.785 | 0.229 |
| Rhythm | Fail | Male | 1.124 | 0.360 |
| Rhythm | Fail | Female | 1.065 | 0.789 |
| Rhythm | Fail | Female | 0.825 | 0.425 |
| Rhythm | Fail | Female | 0.930 | 0.425 |
| Rhythm | Fail | Female | 1.425 | 0.400 |
| Rhythm | Fail | Female | 1.024 | 0.379 |
| Rhythm | Fail | Female | 0.785 | 0.171 |
| Frequency | Success | Male | 0.965 | 0.314 |
| Frequency | Success | Male | 1.308 | 0.760 |
| Frequency | Success | Male | 0.905 | 0.530 |
| Frequency | Success | Male | 0.885 | 0.607 |
| Frequency | Success | Female | 0.805 | 0.469 |
| Frequency | Success | Female | 0.746 | 0.181 |
| Frequency | Success | Female | 1.025 | 0.450 |
| Frequency | Success | Female | 0.745 | 0.260 |
| Frequency | Success | Female | 1.166 | 0.545 |
| Frequency | Success | Female | 0.830 | 0.255 |

## References

Asokan, M. M., Williamson, R. S., Hancock, K. E., & Polley, D. B. (2021). Inverted central auditory hierarchies for encoding local intervals and global temporal patterns. Current Biology, 31(8), 1762-1770.e4. https://doi.org/10.1016/j.cub.2021.01.076

Bååth, R., Tjøstheim, T. A., & Lingonblad, M. (2016). The role of executive control in rhythmic timing at different tempi. Psychonomic Bulletin & Review, 23(6), 1954–1960. https://doi.org/10.3758/s13423-016-1070-1

Benichov, J. I., Benezra, S. E., Vallentin, D., Globerson, E., Long, M. A., & Tchernichovski, O. (2016). The forebrain song system mediates predictive call timing in female and Male Zebra finches. Current Biology, 26(3), 309–318. https://doi.org/10.1016/j.cub.2015.12.037

Berninger, E. (2007). Characteristics of normal newborn transient-evoked otoacoustic emissions: Ear asymmetries and sex effects. International Journal of Audiology, 46(11), 661–669. https://doi.org/10.1080/14992020701438797

Blecher, T., Tal, I., & Ben-Shachar, M. (2016). White matter microstructural properties correlate with sensorimotor synchronization abilities. NeuroImage, 138, 1–12. https://doi.org/10.1016/j.neuroimage.2016.05.022

Bouwer, F. L., Nityananda, V., Rouse, A. A., & ten Cate, C. (2021). Rhythmic abilities in humans and non-human animals: a review and recommendations from a methodological perspective. Philosophical Transactions of the Royal Society B: Biological Sciences, 376(1835). https://doi.org/10.1098/rstb.2020.0335

Brenowitz, E. A., & Remage-Healey, L. (2016). It takes a seasoned bird to be a good listener: Communication between the sexes. Current Opinion in Neurobiology, 38(1), 12–17. https://doi.org/10.1016/j.conb.2016.01.005

Breska, A., & Ivry, R. B. (2018). Double dissociation of single-interval and rhythmic temporal prediction in cerebellar degeneration and Parkinson’s disease. Proceedings of the National Academy of Sciences of the United States of America, 115(48), 12283–12288. https://doi.org/10.1073/pnas.1810596115

Cannon, J. J., & Patel, A. D. (2021). How Beat Perception Co-opts Motor Neurophysiology. Trends in Cognitive Sciences, 25(2), 137–150. https://doi.org/10.1016/j.tics.2020.11.002

Celma-Miralles, A., & Toro, J. M. (2020). Discrimination of temporal regularity in rats (Rattus norvegicus) and humans (Homo sapiens). Journal of Comparative Psychology. https://doi.org/10.1037/com0000202

Chen, J. L., Penhune, V. B., & Zatorre, R. J. (2008). Listening to musical rhythms recruits motor regions of the brain. Cerebral Cortex, 18(12), 2844–2854. https://doi.org/10.1093/cercor/bhn042

Clemens, J., Schöneich, S., Kostarakos, K., Hennig, R. M., & Hedwig, B. (2021). A small, computationally flexible network produces the phenotypic diversity of song recognition in crickets. ELife, 10, 1–31. https://doi.org/10.7554/eLife.61475

Dalla Bella, S., Benoit, C. E., Farrugia, N., Keller, P. E., Obrig, H., Mainka, S., & Kotz, S. A. (2017). Gait improvement via rhythmic stimulation in Parkinson’s disease is linked to rhythmic skills. Scientific Reports, 7(January), 1–11. https://doi.org/10.1038/srep42005

Espinoza-Monroy, M., & de Lafuente, V. (2021). Discrimination of Regular and Irregular Rhythms Explained by a Time Difference Accumulation Model. Neuroscience, 459, 16–26. https://doi.org/10.1016/j.neuroscience.2021.01.035

Gall, M. D., Salameh, T. S., & Lucas, J. R. (2013). Songbird frequency selectivity and temporal resolution vary with sex and season. Proceedings of the Royal Society B: Biological Sciences, 280(1751). https://doi.org/10.1098/rspb.2012.2296

Gámez, J., Mendoza, G., Prado, L., Betancourt, A., & Merchant, H. (2019). The amplitude in periodic neural state trajectories underlies the tempo of rhythmic tapping. PLOS Biology, 17(4), e3000054. https://doi.org/10.1371/journal.pbio.3000054

Gámez, J., Yc, K., Ayala, Y. A., Dotov, D., Prado, L., & Merchant, H. (2018). Predictive rhythmic tapping to isochronous and tempo changing metronomes in the nonhuman primate. Annals of the New York Academy of Sciences, 1–20. https://doi.org/10.1111/nyas.13671

Garcia, M., Theunissen, F., Sèbe, F., Clavel, J., Ravignani, A., Marin-Cudraz, T., Fuchs, J., & Mathevon, N. (2020). Evolution of communication signals and information during species radiation. Nature Communications, 11(1), 1–15. https://doi.org/10.1038/s41467-020-18772-3

Gess, A., Schneider, D. M., Vyas, A., & Woolley, S. M. N. (2011). Automated auditory recognition training and testing. Animal Behaviour, 82(2), 285–293. https://doi.org/10.1016/j.anbehav.2011.05.003

Grahn, J. A., & Brett, M. (2007). Rhythm and beat perception in motor areas of the brain. Journal of Cognitive Neuroscience, 19(5), 893–906. https://doi.org/10.1162/jocn.2007.19.5.893

Grube, M., Cooper, F. E., Chinnery, P. F., & Griffiths, T. D. (2010). Dissociation of duration-based and beat-based auditory timing in cerebellar degeneration. Proceedings of the National Academy of Sciences, 107(25), 11597–11601. https://doi.org/10.1073/pnas.0910473107

Hage, S. R., Gavrilov, N., & Nieder, A. (2016). Developmental changes of cognitive vocal control in monkeys. Journal of Experimental Biology, 219(11), 1744–1749. https://doi.org/10.1242/jeb.137653

Hagmann, C. E., & Cook, R. G. (2010). Testing meter, rhythm, and tempo discriminations in pigeons. Behavioural Processes, 85(2), 99–110. https://doi.org/10.1016/j.beproc.2010.06.015

Honing, H., Bouwer, F. L., Prado, L., & Merchant, H. (2018). Rhesus monkeys (Macaca mulatta) sense isochrony in rhythm, but not the beat: Additional support for the gradual audiomotor evolution hypothesis. Frontiers in Neuroscience, 12, 475. https://doi.org/10.3389/fnins.2018.00475

Hulse, S. H., Humpal, J., & Cynx, J. (1984). Discrimination and Generalization of Rhythmic and Arrhythmic Sound Patterns by European Starlings (Sturnus vulgaris). Music Perception, 1(4), 442–464. https://doi.org/10.2307/40285272

Janik, V. M. (2014). Cetacean vocal learning and communication. Current Opinion in Neurobiology, 28, 60–65. https://doi.org/10.1016/j.conb.2014.06.010

Krentzel, A. A., & Remage-Healey, L. (2015). Sex differences and rapid estrogen signaling: A look at songbird audition. Frontiers in Neuroendocrinology, 38, 37–49. https://doi.org/10.1016/j.yfrne.2015.01.001

Kriengwatana, B., Spierings, M. J., & ten Cate, C. (2016). Auditory discrimination learning in zebra finches: effects of sex, early life conditions and stimulus characteristics. Animal Behaviour, 116, 99–112. https://doi.org/10.1016/j.anbehav.2016.03.028

Krizman, J., Bonacina, S., & Kraus, N. (2020). Sex differences in subcortical auditory processing only partially explain higher prevalence of language disorders in males. Hearing Research, 398, 108075. https://doi.org/10.1016/j.heares.2020.108075

Krizman, J., Rotondo, E. K., Nicol, T., Kraus, N., & Bieszczad, K. (2021). Sex differences in auditory processing vary across estrous cycle. Scientific Reports, 11(1), 1–7. https://doi.org/10.1038/s41598-021-02272-5

Krotinger, A., & Loui, P. (2021). Rhythm and groove as cognitive mechanisms of dance intervention in Parkinson’s disease. PLoS ONE, 16(5 May), 1–20. https://doi.org/10.1371/journal.pone.0249933

Levitin, D. J., & Cook, P. R. (1996). Memory for musical tempo: Additional evidence that auditory memory is absolute. Perception and Psychophysics, 58(6), 927–935. https://doi.org/10.3758/BF03205494

Mathevon, N., Casey, C., Reichmuth, C., & Charrier, I. (2017). Northern Elephant Seals Memorize the Rhythm and Timbre of Their Rivals’ Voices. Current Biology, 27(15), 2352-2356.e2. https://doi.org/10.1016/j.cub.2017.06.035

Merchant, H., Pérez, O., Bartolo, R., Méndez, J. C., Mendoza, G., Gámez, J., Yc, K., & Prado, L. (2015). Sensorimotor neural dynamics during isochronous tapping in the medial premotor cortex of the macaque. European Journal of Neuroscience, 41(5), 586–602. https://doi.org/10.1111/ejn.12811

Merker, B. H., Madison, G. S., & Eckerdal, P. (2009). On the role and origin of isochrony in human rhythmic entrainment. Cortex, 45(1), 4–17. https://doi.org/10.1016/j.cortex.2008.06.011

Nagel, K. I., McLendon, H. M., & Doupe, A. J. (2010). Differential Influence of Frequency, Timing, and Intensity Cues in a Complex Acoustic Categorization Task. Journal of Neurophysiology, 104(3), 1426–1437. https://doi.org/10.1152/jn.00028.2010

Naqvi, S., Godfrey, A. K., Hughes, J. F., Goodheart, M. L., Mitchell, R. N., & Page, D. C. (2019). Conservation, acquisition, and functional impact of sex-biased gene expression in mammals. Science, 365(6450). https://doi.org/10.1126/science.aaw7317

Norton, P., & Scharff, C. (2016). ‘Bird song metronomics’: Isochronous organization of zebra finch song rhythm. Frontiers in Neuroscience, 10(JUL), 97–99. https://doi.org/10.3389/fnins.2016.00309

Nottebohm, F., & Arnold, A. P. (1976). Sexual Dimorphism in Vocal Control Areas of the Songbird Brain. Science, 194(4261), 211–213. https://doi.org/10.1126/science.959852

Patel, A. D. (2006). Musical rhythm, linguistic rhythm, and human evolution. Music Perception, 24(1), 99–103. https://doi.org/10.1525/Mp.2006.24.1.99

Patel, A. D., & Iversen, J. R. (2014). The evolutionary neuroscience of musical beat perception: the Action Simulation for Auditory Prediction (ASAP) hypothesis. Frontiers in Systems Neuroscience, 8(May), 57. https://doi.org/10.3389/fnsys.2014.00057

Rajendran, V. G., Harper, N. S., Garcia-Lazaro, J. A., Lesica, N. A., & Schnupp, J. W. H. (2017). Midbrain adaptation may set the stage for the perception of musical beat. Proceedings of the Royal Society B: Biological Sciences, 284(1866). https://doi.org/10.1098/rspb.2017.1455

Ravignani, A., & Madison, G. (2017). The Paradox of Isochrony in the Evolution of Human Rhythm. Frontiers in Psychology, 8. https://doi.org/10.3389/fpsyg.2017.01820

Roeske, T. C., Tchernichovski, O., Poeppel, D., & Jacoby, N. (2020). Categorical Rhythms Are Shared between Songbirds and Humans. Current Biology, 30(18), 3544-3555.e6. https://doi.org/10.1016/j.cub.2020.06.072

Ross, J. M., Iversen, J. R., & Balasubramaniam, R. (2018). The Role of Posterior Parietal Cortex in Beat-based Timing Perception: A Continuous Theta Burst Stimulation Study. Journal of Cognitive Neuroscience, 30(5), 634–643. https://doi.org/10.1162/jocn_a_01237

Rouse, A. A., Patel, A. D., & Kao, M. H. (2021). Vocal learning and flexible rhythm pattern perception are linked: Evidence from songbirds. Proceedings of the National Academy of Sciences, 118(29), e2026130118. https://doi.org/10.1073/pnas.2026130118

Samuels, B., Grahn, J., Henry, M. J., & Macdougall-Shackleton, S. A. (2021). European starlings (sturnus vulgaris) discriminate rhythms by rate, not temporal patterns. The Journal of the Acoustical Society of America, 149, 2546. https://doi.org/10.1121/10.0004215

Savage, P. E., Brown, S., Sakai, E., & Currie, T. E. (2015). Statistical universals reveal the structures and functions of human music. Proceedings of the National Academy of Sciences of the United States of America, 112(29), 8987–8992. https://doi.org/10.1073/pnas.1414495112

Schiavo, J. K., Valtcheva, S., Bair-Marshall, C. J., Song, S. C., Martin, K. A., & Froemke, R. C. (2020). Innate and plastic mechanisms for maternal behaviour in auditory cortex. Nature, 587(7834), 426–431. https://doi.org/10.1038/s41586-020-2807-6

Schöneich, S., Kostarakos, K., & Hedwig, B. (2015). An auditory feature detection circuit for sound pattern recognition. Science Advances, 1(8). https://doi.org/10.1126/sciadv.1500325

Shaughnessy, D. W., Hyson, R. L., Bertram, R., Wu, W., & Johnson, F. (2019). Female zebra finches do not sing yet share neural pathways necessary for singing in males. Journal of Comparative Neurology, 527(4), 843–855. https://doi.org/10.1002/cne.24569

Stansbury, A. L., & Janik, V. M. (2019). Formant Modification through Vocal Production Learning in Gray Seals. Current Biology. https://doi.org/10.1016/j.cub.2019.05.071

Takahashi, D. Y., Liao, D. A., & Ghazanfar, A. A. (2017). Vocal Learning via Social Reinforcement by Infant Marmoset Monkeys. Current Biology, 27(12), 1844-1852.e6. https://doi.org/10.1016/j.cub.2017.05.004

Takeya, R., Kameda, M., Patel, A. D., & Tanaka, M. (2017). Predictive and tempo-flexible synchronization to a visual metronome in monkeys. Scientific Reports, 7(1), 1–12. https://doi.org/10.1038/s41598-017-06417-3

Teki, S., Grube, M., Kumar, S., & Griffiths, T. D. (2011). Distinct Neural Substrates of Duration-Based and Beat-Based Auditory Timing. Journal of Neuroscience, 31(10), 3805–3812. https://doi.org/10.1523/JNEUROSCI.5561-10.2011

Ten Cate, C., Spierings, M., Hubert, J., & Honing, H. (2016). Can birds perceive rhythmic patterns? A review and experiments on a songbird and a parrot species. Frontiers in Psychology, 7(MAY). https://doi.org/10.3389/fpsyg.2016.00730

Trehub, S. E., & Thorpe, L. A. (1989). Infants’ perception of rhythm: categorization of auditory sequences by temporal structure. Canadian Journal of Psychology, 43(2), 217–229. https://doi.org/10.1037/h0084223

van der Aa, J., Honing, H., & ten Cate, C. (2015). The perception of regularity in an isochronous stimulus in zebra finches (Taeniopygia guttata) and humans. Behavioural Processes, 115, 37–45. https://doi.org/10.1016/j.beproc.2015.02.018

Vaquero, L., Ramos-Escobar, N., François, C., Penhune, V., & Rodríguez-Fornells, A. (2018). White-matter structural connectivity predicts short-term melody and rhythm learning in non-musicians. NeuroImage, 181, 252–262. https://doi.org/10.1016/j.neuroimage.2018.06.054

Verga, L., Sroka, M. G. U., Varola, M., Villanueva, S., & Ravignani, A. (2022). Spontaneous rhythm discrimination in a mammalian vocal learner. Biology Letters, 18(10). https://doi.org/10.1098/rsbl.2022.0316

Vernes, S. C., & Wilkinson, G. S. (2019). Behaviour, biology and evolution of vocal learning in bats. Philosophical Transactions of the Royal Society B: Biological Sciences, 375(1789), 20190061. https://doi.org/10.1098/rstb.2019.0061

Wirthlin, M., Chang, E. F., Knörnschild, M., Krubitzer, L. A., Mello, C. v, Miller, C. T., Pfenning, A. R., Vernes, S. C., Tchernichovski, O., & Yartsev, M. M. (2019). A Modular Approach to Vocal Learning: Disentangling the Diversity of a Complex Behavioral Trait. Neuron, 104(1), 87–99. https://doi.org/10.1016/j.neuron.2019.09.036

Yeh, Y.-T., Rivera, M., & Woolley, S. M. N. (2022). Auditory sensitivity and vocal acoustics in five species of estrildid songbirds. Animal Behaviour. https://doi.org/10.1016/j.anbehav.2022.11.002

Zann, R. A. (1996). The zebra finch: a synthesis of field and laboratory studies. Oxford University Press.

Zentner, M. (2021). Social bonding and credible signaling hypotheses largely disregard the gap between animal vocalizations and human music. Behavioral and Brain Sciences, 44, e120. https://doi.org/10.1017/S0140525X2000165X

